# Night light of a metropolitan coastline aligns with habitat segregation and divergent plasticity among closely related coastal isopods

**DOI:** 10.1101/2025.10.22.684051

**Authors:** Daiki X. Sato

## Abstract

Artificial light at night (ALAN) is a pervasive urban driver that can reshape coastal communities, yet its links to genetic structure and behavior remain unclear. Here, we examined two closely related isopods, *Ligia laticarpa* and *L. furcata*, across Tokyo Bay along a gradient from brightly lit inner-bay shorelines to darker, vegetated outer-bay coasts. Genomic and mitochondrial evidence consistently identified clear genetic segregation between outer- and inner-bay populations corresponding to the two species, with *L. laticarpa* predominating at inner-bay and *L. furcata* at outer-bay sites. While individual genetic profiles showed little indication of recent hybridization, population signals in several inner-bay sites pointed to historical inputs or local co-occurrence of distinct lineages, possibly mediated by ships. Bayesian modeling of 28 years of environmental data identified nighttime light intensity, salinity, and vegetation cover as major predictors of species occurrence, highlighting ALAN as a key environmental axis distinguishing their habitats. In laboratory assays, chronic ALAN exposure reduced growth and activity in *L. furcata* but had limited effects on *L. laticarpa*. These findings suggest that artificial lighting and maritime activity jointly shape patterns of genetic connectivity, habitat segregation, and physiological plasticity between closely related coastal species, providing insights into how anthropogenic disturbances influence ecological divergence.

## Introduction

Urbanization is a pervasive global process that reshapes natural environments and imposes novel selective pressures on organisms. Among the many anthropogenic disturbances associated with urban development, artificial light at night (ALAN) and increased human mobility can profoundly affect animal populations by altering their physiology, behavior, and spatial distributions. ALAN disrupts natural light–dark cycles and influences circadian rhythms, predator–prey dynamics, and reproduction across a wide range of taxa (1–7). For nocturnal or crepuscular species, whose activity patterns are tightly regulated by natural light cycles, ALAN may be a particularly strong driver of ecological divergence and evolutionary change, potentially altering species persistence and distribution (8–10). Yet, how such sensory pollution interacts with other human disturbances to shape ecological and genetic patterns in closely related species remains largely unexplored.

Coastal ecosystems are among the most strongly affected by urbanization, with intense nighttime illumination and high levels of human traffic (11–13). Crustaceans and other intertidal organisms are particularly sensitive to changes in light regimes (14, 15): barnacles and crabs show altered settlement, emergence, and survival under artificial illumination (16, 17), and semi-terrestrial amphipods and isopods exhibit changes in locomotion and circadian gene expression (18–21). The common sea slater (*Ligia oceanica*) has been reported to exhibit behavioral and morphological changes linked to anti-predator responses under ALAN (22). These findings underscore the profound influence of artificial lighting on coastal ecosystems and highlight how even modest changes in nighttime illumination can affect a range of behavioral and physiological traits. Nevertheless, many aspects of ALAN’s broader ecological and evolutionary consequences in these habitats remain poorly understood.

The coastal isopods of the genus *Ligia* provide a unique system for studying such processes. Two closely related species, *L. furcata* and *L. laticarpa*, were recently distinguished based on morphological and genetic differences (23). Both inhabit the rocky intertidal zones along Tokyo Bay, a megacity shoreline that is heavily engineered and trafficked, and ranks among the most intensely illuminated coastal regions worldwide.

Despite their limited natural dispersal ability, *Ligia* species are frequently found near ports and artificial shorelines, raising the possibility that human-mediated environmental changes and/or transport may influence their population divergence and connectivity. In addition, *Ligia* is primarily nocturnal (24) and highly sensitive to light (22), implying that artificial illumination could alter their physiology and habitat use in species-specific ways.

In this study, we combined field genomic, environmental, and experimental approaches to examine how anthropogenic disturbances relate to genetic and ecological differentiation in *Ligia*. We first characterized species composition, population structure, and admixture with target sequencing and a genome-wide SNP analysis across populations along the coastline of Tokyo Bay. We then identified environmental variables underlying species distributions using long-term remote-sensing data and Bayesian modeling. Finally, we examined species-specific physiological and behavioral responses to chronic artificial light exposure under controlled laboratory conditions. This combination of approaches allows us to link population genetic structure, environmental gradients, and phenotypic plasticity, providing new insight into how urban disturbances shape genetic and ecological boundaries in coastal organisms.

## Results

### Genetic and environmental differentiation between inner- and outer-bay populations

We first sequenced mitochondrial 16S and nuclear 18S rRNA genes from 26 populations along the Tokyo Bay coastline and a population from Hokkaido as an outgroup (see Table S1 for detailed sampling information). Based on comparisons with previous studies (23), we confirmed the presence of three species across Tokyo Bay populations: *L. furcata* (outer bay), *L. laticarpa* (inner bay), and *L. cinerascens* (occasionally at inner bay) (Fig. 1a; Fig. S1). We then analyzed genome-wide SNPs from MIG-seq; after filtering, 193 SNPs were retained. ADMIXTURE analyses for *K* = 2–8 showed a marked decrease in cross-validation (CV) error up to *K* = 3, with a slight further reduction at *K* = 4 (Fig. S2). The additional cluster at *K* = 4 represented a subdivision within *L. furcata* populations, corresponding to the two regional clades. Because *K* = 3 best reflected the three major species detected in our dataset, we present results for *K* = 3 in the main text. Together with principal component analysis (PCA), the ADMIXTURE result recovered two dominant clusters that matched *L. laticarpa* in inner-bay sites and *L. furcata* on outer-bay coasts, with occasional *L. cinerascens* occurring at inner-bay sites (Fig. 1b). At the individual level, most samples showed near-pure ancestry within a single cluster, indicating little evidence of recent hybrid individuals in our dataset at our marker density. We next evaluated population-level admixture signals using three-population *f3* statistics. Negative *f*_*3*_ values in a target population indicate that its allele-frequency covariance can be explained by contributions from two source groups. Importantly, this does not imply that sampled individuals are hybrids; rather, it shows that the target population as a whole carries ancestry components related to the sources, which can arise from historical gene flow, secondary contact, or co-occurrence of distinct lineages at the same site. A subset of inner-bay populations showed *f*_*3*_ patterns consistent with contributions from lineages related to nearby groups, including *L. cinerascens*. These population-level migration signals are compatible with the individual-level ADMIXTURE result of largely non-admixed genotypes, and together they suggest local co-occurrence or historical inputs rather than pervasive, ongoing hybridization. Given the moderate number of SNPs, these results should be interpreted as broad-scale signals of population structure rather than fine-scale admixture. An association with local vessel density (mean monthly number of AIS-equipped vessels within 1 km of each site) possibly suggests their ship-mediated dispersal among coastal sites (Fig. 1c).

**Figure 1.**
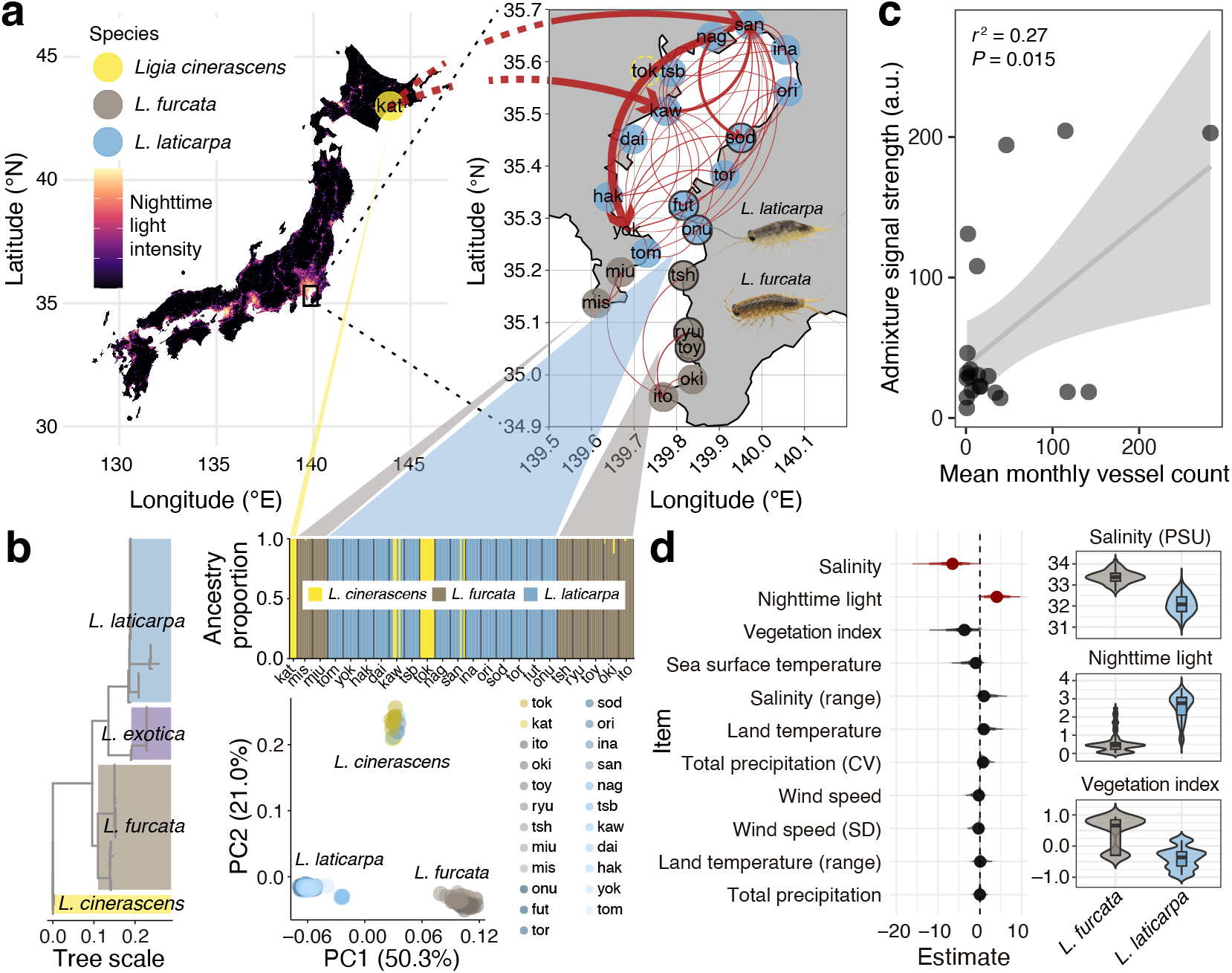
Geographic distribution, genetic structure, and environmental differentiation of *Ligia* species along the Tokyo Bay coastline. **(a)** Sampling sites of *L. cinerascens* (yellow), *L. furcata* (brown), and *L. laticarpa* (blue) across Japan, overlaid on nighttime light intensity (radiance, log-scaled) derived from the SRUNet_NPP_VIIRS_V2_Like_2020 dataset. An enlarged view highlights the Tokyo Bay coastline. Red arrows depict population-level estimates of migration directions and magnitudes based on *f*_*3*_ statistics. Given the plausibly complex admixture history, a population from Hokkaido (kat), rather than the inner Tokyo Bay population (tok), was used as an outgroup representing *L. cinerascens*. Photo credit: author. **(b)** (Left) Maximum-likelihood tree of 16S rRNA gene sequences from 27 populations investigated in this study and reference sequences (see Fig. S1 for details). (Top right) Ancestry proportions estimated by ADMIXTURE (*K* = 3), corresponding to the three species identified by mitochondrial 16S sequencing. (Bottom right) Principal component analysis (PCA) showing clear separation between the three species. **(c)** Positive correlation between admixture signal strength (*f*_*3*_ Z-scores) and mean monthly vessel density within 1 km of each sampling site (*r*^*2*^ = 0.27, *P* = 0.015), suggesting a potential role of ship-mediated dispersal. (**d**) (Left) Bayesian modeling of 28 years of remotely sensed environmental variables identified salinity, nighttime light intensity, and vegetation index as the strongest predictors of *L. furcata* and *L. laticarpa* occurrence (see Table S2 for details). Response coded as *L. laticarpa* = 1, and positive coefficients indicate higher probability of *L. laticarpa* occurrence. Variables with *p*_*d*_ > 0.95 are colored in red. (Right) Violin plots show corresponding environmental differences between the habitats of the two species.

Long-term environmental data analyses further revealed pronounced ecological differentiation between the habitats of two species. Bayesian modeling of 28 years of remotely sensed climate and oceanographic variables identified nighttime light intensity, salinity, and vegetation cover as the strongest predictors of species occurrence. *Ligia furcata* occurrence was associated with higher salinity (posterior mean = −7.01, 95% CI = −16.16 to −0.25, *p*_*d*_ = 0.98) and denser, more vegetated habitats (−4.12, −12.12 to 0.16, 0.94), whereas *L. laticarpa* occurrence increased with higher nighttime light intensity (4.15, −0.03 to 9.74, 0.97) (Figs. 1d and S3; Table S2).

### Chronic ALAN exposure influences the survival and growth of *Ligia*

To examine the effect of chronic ALAN exposure on the physiological states of these species, we collected samples from three populations located near the boundary of their respective habitats (toy, ryu, and tsh populations for *L. furcata* and onu, fut, and sod populations for *L. laticarpa*; highlighted with black-rimmed circles in Fig. 1a) and reared them for five months under either ALAN or control conditions, as shown in Fig. S4. Survival probability was significantly influenced by both effects of species and ALAN: Chronic ALAN exposure was associated with higher survival in both species (*χ*^2^ = 22.75, df = 1, *P* < 0.001, Cox proportional hazards regression model), whereas *L. furcata* and *L. laticarpa* differed markedly in their overall survival rates (*χ*^2^ = 27.67, df = 1, *P* < 0.001, Cox proportional hazards regression model) (Fig. 2a). In addition, body length measurements revealed no significant main effect of ALAN exposure on growth (*χ*^2^ = 1.622, df = 1, *P* = 0.203, LMM), nor a main effect of species (*χ*^2^ = 0.778, df = 1, *P* = 0.378). However, we observed a significant interaction between species and rearing condition (*χ*^2^ = 5.178, df = 1, *P* = 0.023), suggesting that the effect of ALAN on growth differed between species. Furthermore, there were strong interactions between species and rearing time (*χ*^2^ = 34.135, df = 2, *P* = 5.14 × 10^−9^), and between rearing condition and time (*χ*^2^ = 7.904, df = 2, *P* = 0.0049), indicating that growth trajectories over time were influenced by both effects of species and ALAN exposure (Table S3, Fig. 2b).

**Figure 2.**
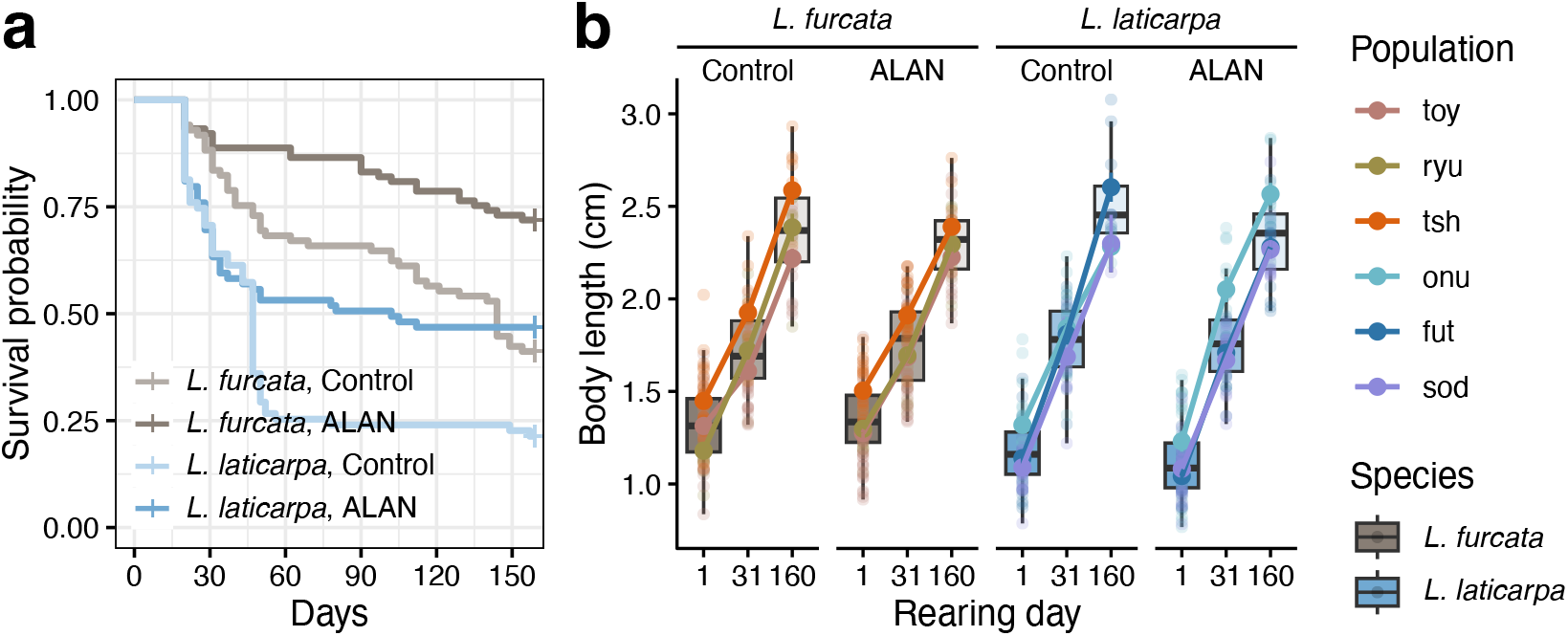
Effects of chronic ALAN exposure on survival and growth. **(a)** Kaplan–Meier survival curves for both species under ALAN and control conditions. **(b)** Body length (cm) measured at three time points (day 1, 31, and 160) under each condition. Boxplots and coloured lines represent population means, with error bars indicating standard errors. Statistical details analyzed with linear mixed models are described in Table S3.

### ALAN differentially affects the activity rhythms of the two species

Individuals reared for five months were then monitored for locomotor activity under a 12L:12D cycle (with the dark period maintained under ALAN or control conditions) over the course of one day. A marked increase in activity at the onset of morning light was observed in many individuals (Fig. 3a), with statistical significance confirmed in both species under most conditions (Fig. 3b; \*\*\**P* < 0.001, \*\**P* < 0.01, \**P* < 0.05, Wilcoxon’s signed rank test). However, this morning activity peak was absent in individuals that had not been reared under ALAN but were only exposed to it acutely during the experiment. Furthermore, we found that chronic exposure to ALAN had a significant effect on mean daily activity levels (*χ*^2^ = 4.519, df = 1, *P* = 0.034), while the light condition during the behavioral test itself had no significant effect (*χ*^2^ = 0.996, df = 1, *P* = 0.318; Table S4). The two species also differed significantly in their overall activity (*χ*^2^ = 4.498, df = 1, *P* = 0.034), and a significant interaction between species and rearing condition (*χ*^2^ = 7.231, df = 1, *P* = 0.007) indicated that the effect of chronic ALAN exposure varied between them. Specifically, *L. furcata* showed reduced activity when reared under ALAN, while *L. laticarpa* was largely unaffected (Fig. 3c). These results suggest that it is the long-term effects of ALAN during development, rather than the immediate lighting environment during testing, that drive the observed species-specific differences in activity rhythms.

**Figure 3.**
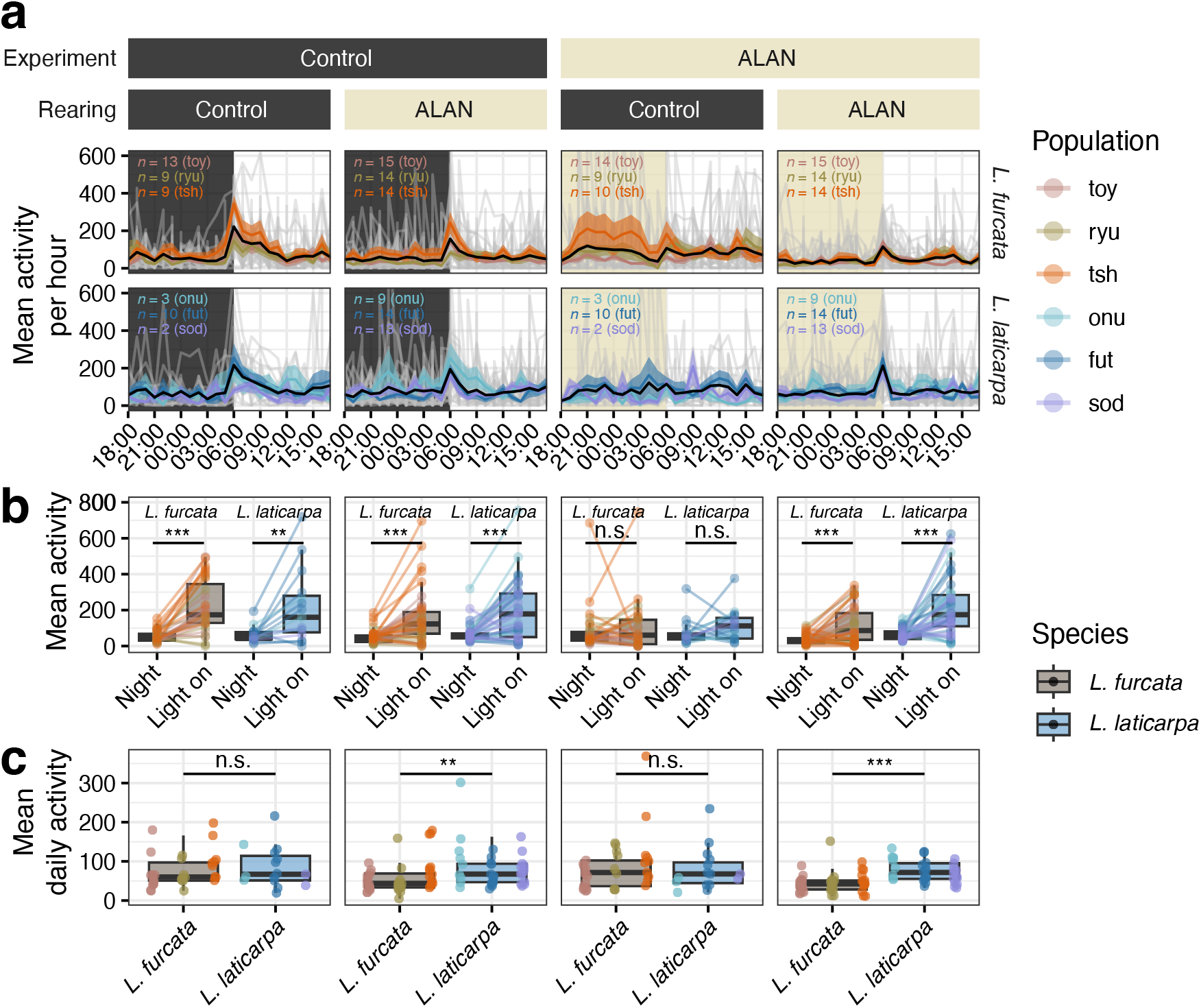
Effects of chronic ALAN exposure on daily locomotor activity in *Ligia furcata* and *L. laticarpa*. **(a)** Mean activity per hour profiles under two rearing conditions (control or ALAN) and two experimental conditions (control or ALAN). The grey and black lines indicate individual values and the overall mean value of activity, respectively. Each coloured line denotes the mean value for a different population, with corresponding sample sizes indicated. Shaded areas around the lines represent the standard error of the mean. **(b)** Box plots comparing mean activity levels during the night (‘Night’) and the light-on period (‘Light on’) for each combination of rearing and experimental condition. Asterisks indicate significant differences between periods, evaluated using Wilcoxon’s signed rank test (\*\*\**P* < 0.001, \*\**P* < 0.01, \**P* < 0.05); n.s., not significant. **(c)** Mean 24-hour activity of *Ligia furcata* and *L. laticarpa* under each rearing condition (control vs. ALAN). Asterisks mark significant interspecific differences, evaluated with Wilcoxon’s rank sum test. Statistical details are described in Table S4.

Finally, to assess whether the observed activity patterns reflected endogenous circadian rhythms, locomotor activity was recorded continuously for four days under complete darkness. Under these dark conditions, overall rhythmicity was weak at the group level, yet period estimates from individual traces showed opposite tendencies between species (Figs. 4a and S5). A LMM analysis revealed a significant interaction between species and rearing condition (*χ*^2^ = 4.24, df = 1, *P* = 0.039; Table S5), indicating that the effect of ALAN exposure during development differed between species. Specifically, *L. laticarpa* showed a trend toward increased activity, whereas *L. furcata* exhibited a decrease in activity following chronic ALAN exposure (Fig. 4b).

**Figure 4.**
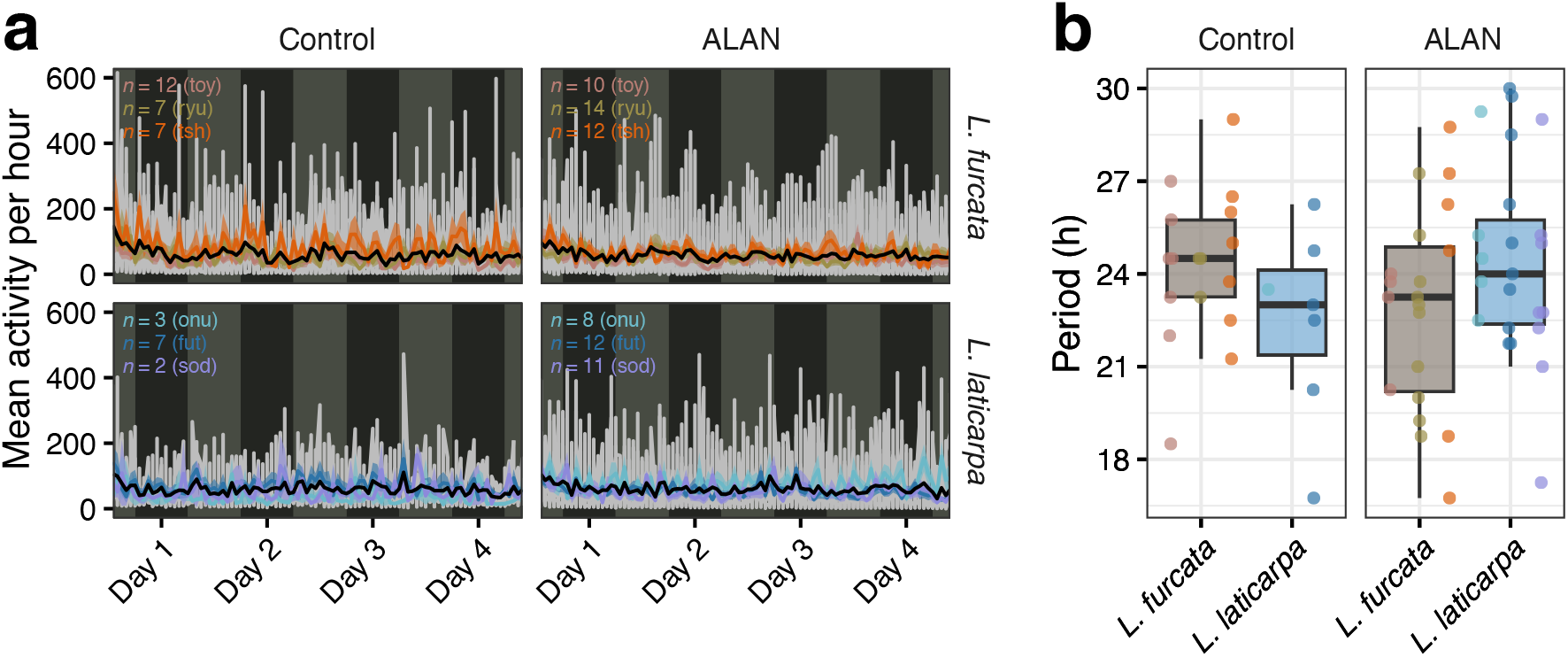
Endogenous rhythms of *Ligia furcata* and *L. laticarpa* under continuous darkness. **(a)** Mean locomotor activity profiles recorded over four consecutive days in constant darkness, comparing individuals reared under control (left panels) and ALAN (right panels) conditions. Grey and black lines indicate individual values and the overall mean activity, respectively. Each coloured line represents a different population, with corresponding sample sizes indicated. Shaded areas around the lines represent the standard error of the mean. Brighter and darker backgrounds indicate daytime and nighttime, respectively. **(b)** Box plots of circadian period (in hours), estimated by periodogram analysis, for control- and ALAN-reared individuals of each species. Points represent individual values. Statistical details are provided in Table S5.

## Discussion

Our study clarifies the broad biogeographic context of closely related *Ligia* in Japan by integrating mitochondrial and nuclear markers with prior taxonomic work (23) and the nationwide COI-based surveys (25). Based on the correspondence between our sampling localities and the COI clades, *L. exotica* Groups E (Izu) and F (Hachijo) (25) likely correspond to *L. furcata*, whereas Groups A–C and possibly D (Lineage I-1) correspond to *L. laticarpa*. This synthesis, together with the nationwide distribution map (Fig. S6), indicates that while *L. furcata* is confined to the Kanto and southern Tohoku regions, *L. laticarpa* occupies much of the Japanese coastline, particularly in western areas. However, within the Kanto region, *L. laticarpa* is restricted to inner-bay shorelines of Tokyo Bay, surrounded by *L. furcata* populations. Integrating Itani’s proposal (25), *L. furcata*, originally distributed in the Izu Islands, likely expanded northward onto Honshu during coastline reconfiguration associated with the development of the Izu–Bonin arc (26) and the formation of the Izu Peninsula. It remains unclear whether the restricted distribution of *L. laticarpa* in the Kanto region results from ecological competition with the expanding *L. furcata* or other locally co-occurring *Ligia* species (*L. exotica* and *L. cinerascens*), or from secondary introduction from western Japan through human-mediated transport. Comprehensive genomic analyses with broader sampling across Japan will be necessary to test these alternative scenarios and clarify the evolutionary processes shaping their present distributions.

Within Tokyo Bay, species composition and spatial arrangement are clearly delineated. *Ligia laticarpa* predominates at inner-bay shorelines, whereas *L. furcata* is more common along outer-bay shorelines. Individual genotypes were largely non-admixed, and we did not detect any apparent hybrids in our sampling. At the population level, several inner-bay populations showed signals consistent with the local co-occurrence of *L. laticarpa* and *L. cinerascens*. Considering that the major range of *L. cinerascens* along the Japanese coast is in northern regions such as Hokkaido and Tohoku (Fig. S6), these observations are most consistent with secondary presence in Tokyo Bay facilitated by human activity, plausibly ship-mediated transport. The distributional patterns of *L. laticarpa* and *L. furcata* align with their significant environmental contrasts between inner and outer bay, notably nighttime light, salinity, and shoreline vegetation. This setting provides a natural context for evaluating how closely related species partition coastal habitats under urban light regimes.

Our rearing experiments revealed physiological effects of chronic ALAN exposure, particularly on survival and growth. Survival probability was higher under ALAN in both species (Fig. 2a). Because mortality often coincided with moulting, group housing may have amplified moulting-related predation reported in crustaceans (27). A plausible explanation is that artificial light increased visual detectability of conspecific movements, allowing individuals to better avoid encounters that could lead to opportunistic cannibalism. In this way, ALAN may have indirectly reduced molting-related mortality and increased apparent survival. However, because individuals were reared in groups, we cannot exclude other indirect effects through altered social interactions or activity timing, and direct physiological benefits cannot be inferred from our design. Both species tended to show reduced growth under ALAN, with a smaller reduction in *L. laticarpa* than in *L. furcata* (Fig. 2b and Table S3). Growth trajectories over time were also influenced by both rearing condition and species, indicating that developmental responses to nighttime light exposure diverge between the two species. Growth in isopods can also be modulated by social interactions, such as pheromone-mediated synchronization of molting or reduced desiccation stress through aggregation (28). Individually housed, longitudinal trials will therefore be needed to separate ALAN effects from social context.

Chronic ALAN exposure had species-specific effects on activity rhythms. A lights-on surge in locomotion was generally observed (Fig. 3a), but it disappeared when ALAN was applied only during the test to animals reared under control conditions (Figs. 3a–b). This suggests that the prior light environment during rearing has a lasting effect on the lights-on response. Mean daily activity differed between species and depended on rearing condition (Fig. 3c), with body size as an important covariate (Table S4). Under constant darkness, prior ALAN exposure appeared to shift endogenous periods in opposite directions—lengthening in *L. laticarpa* and shortening in *L. furcata* (Fig. 4b). This divergence suggests species-specific plasticity of the circadian oscillator to developmental ALAN; whether these opposite shifts confer any functional advantage in brighter inner-bay habitats will require targeted performance assays. Although *Ligia* species are generally regarded as nocturnal, such a pattern was not clearly observed in this study. The pronounced responsiveness to morning light, on the other hand, is consistent with strong activity around dawn (29). Nevertheless, behavioral rhythms measured in the laboratory may differ from those in the field, and as shown here, long-term rearing under artificial conditions can itself modify activity timing. Further work is therefore needed to characterize the natural circadian rhythm of these species.

Our results show that two closely related coastal isopods sort along an urban light gradient, with *L. laticarpa* predominating in brighter, inner-bay shorelines and *L. furcata* in darker, outer-bay coasts. At the individual level we found little evidence of recent hybrids, while population-level signals indicate that human activity, especially ship traffic, can promote local co-occurrence of distinct lineages (i.e., *L. cinerascens*) without obvious genetic blending. Together with laboratory rearing, which demonstrates durable, species-specific effects of developmental ALAN on activity rhythms, these findings point to ecological partitioning under modern light regimes and to evolved or plastic differences in light sensitivity that help maintain species boundaries in close contact. More broadly, the present study adds to a growing view that contemporary human disturbance can shape evolution not only by imposing selection but also by altering contact zones and dispersal pathways. The species that occupies more urbanized shores has a wider geographic footprint and shows responses consistent with greater phenotypic flexibility, whereas the species of less-lit coasts appears narrower in range and more sensitive to experimental light histories. This contrast raises a general hypothesis: habitat breadth in human-altered environments may track the capacity for developmental plasticity in sensory and circadian systems. Such interspecific variation in ALAN vulnerability and light-intensity thresholds has been recently reported across birds (6, 7), suggesting that tolerance breadth to illuminated environments may similarly depend on species-specific plasticity in light sensitivity.

Addressing this hypothesis will require integrative approaches, combining denser genomic data, controlled rearing experiments, and manipulative field work to disentangle developmental from acute light effects. Furthermore, although our study focused on Tokyo Bay to identify environmental differentiation and plastic behavioral responses between the two species, future nation- or continent-wide genomic and ecological surveys will be essential to uncover the evolutionary foundations of environmental adaptation in *Ligia*. Pursuing these steps will strengthen causal links between urban light, habitat partitioning, and the evolution of plasticity, and will inform light-management strategies for biodiversity in rapidly developing coastlines.

## Materials and methods Ethical statement

All sampling and rearing procedures followed institutional and national guidelines for the ethical use of invertebrates in research and were approved by the Ethics Committee for Animal Experiments at Chiba University (DOU29-203).

### Field sampling and species identification via target sequencing

A total of 26 populations of *Ligia* were comprehensively sampled from the coastline of Tokyo Bay (eight individuals per population), along with one population from Kushiro, Hokkaido (four individuals). Sample information is summarized in Table S1. For preliminary species identification, we examined mitochondrial 16S rRNA and nuclear 18S rRNA gene sequences using the following protocol. For each population, mitochondrial and nuclear DNA were extracted from pleopods of two individuals using MightyPrep reagent (#9182, Takara Bio Inc., Japan) and amplified via PCR with MightyAmp DNA Polymerase (#R076A, Takara Bio Inc.). The mitochondrial 16S fragment was amplified using primers 16Sar-L (5’-CGCCTGTTTATCAAAAACAT-3’) and 16Sbr-H (5’-CCGGTCTGAACTCAGATCACGT-3’) (30), and the nuclear 18S fragment using primers LoV7F (5’-GGGACCACCAGGAGTG-3’) and LOV7R (5’-GGCCCAGAACATCTAAGG-3’) (31). The thermal cycling conditions consisted of an initial denaturation at 98 °C for 2 min, followed by 30 cycles of 98 °C for 10 s, 58 °C for 30 s and 68 °C for 1 min. Sanger-sequenced 16S or 18S rRNA gene sequences were aligned using the mafft v7.525 (32) with ‘-auto’ option and compared with the previously sequenced *Ligia* samples (23, 33–38). Based on the nucleotide differences observed in the alignment (see Supplementary Data 1), *L. laticarpa* was identified by a substitution at position 64 (G; previously reported as 63G (23)), whereas *L. furcata* possessed 244C. We also detected diagnostic substitutions for *L. exotica* and *L. cinerascens* at positions 177G (previously 175G (23)) and 367C, respectively (Fig. S1c). These species-specific markers enabled confident assignment of individuals and populations to each species.

### Population genetic analyses with MIG-seq

Genomic DNA was extracted from samples of 23 out of the 27 populations surveyed (Fig. 1a; see Table S1) using the Maxwell® RSC Plant DNA Kit (AS1490, Promega, Madison, WI, USA). PCR-amplified DNA was then subjected to multiplexed ISSR genotyping by sequencing (MIG-seq) (39) to infer fine-scale genetic structure and species composition across sites. Library preparations and sequencing with an Illumina MiSeq were performed at GENODAS (Miyagi, Japan). Raw paired-end reads were quality-filtered and trimmed using fastp and cutadapt to remove low-quality bases and standardize read length to 80 bp.

Processed reads were analyzed with the Stacks pipeline (v2.62) in de novo mode to assemble loci and call SNPs. Parameter settings included a minimum stack depth of 3, within-individual mismatch threshold of 2, between-individual mismatch threshold of 2, minimum minor allele frequency of 0.05, maximum observed heterozygosity of 0.60, and inclusion of loci present in at least 70% of individuals in at least 18 populations.

After filtering, 193 SNPs were retained for population structure analyses. ADMIXTURE (v1.3.0) (40) was used to estimate ancestry components, with cross-validation performed for *K* values ranging from 2 to 8 to determine the optimal number of clusters (Fig. S2). Principal component analysis (PCA) based on the same SNP set was conducted using PLINK to visualize genetic differentiation among populations. To further evaluate historical relationships and potential gene flow, TreeMix (41) and *f*_*3*_ statistics were applied. Given the likely complex admixture history, a population identified as *L. cinerascens* within Tokyo Bay (tok) was excluded from these analyses, while the *L. cinerascens* population from Kushiro was used as a representative outgroup to infer migration events involving this species. Z-scores from the *f*_*3*_ test were then used to visualize the relative strength and direction of inferred migration among regional populations, and for clarity, only the top 50 migration edges (with the strongest negative Z-scores) were shown in the final network visualization.

### Quantification of human maritime activity

To quantify the level of human maritime activity around each population site, we obtained vessel tracking data from the Global Fishing Watch (GFW) database (https://globalfishingwatch.org). Using the GFW API, we retrieved all records of AIS-equipped vessels detected within a 1 km radius of each sampling location over the past five years (January 2020 to September 2025). For each population, the total number of unique vessels present within this radius was aggregated by month, and the mean monthly vessel count was calculated as an index of local maritime traffic intensity. This measure was used to represent the long-term average exposure of each population to anthropogenic disturbance associated with coastal vessel activity and was also used to test for correlations with the degree of genetic admixture among populations.

### Environmental data collection and analysis

To compare environmental conditions across habitats of *L. furcata* and *L. laticarpa*, we compiled terrestrial and marine variables from publicly available remote-sensing products. Terrestrial climate and vegetation were obtained from the Copernicus Climate Data Store (https://cds.climate.copernicus.eu; ERA5-Land monthly means), and marine conditions from the Copernicus Marine Service (https://marine.copernicus.eu; Global Ocean Physics Reanalysis, ORAS5). For each dataset, we assembled a multi-year time series spanning January 1993 to December 2020 (using the longest available record within this window for each product). All rasters were projected to WGS84 and values were extracted at each population site; for marine variables at coastal sites, values were taken from the nearest valid ocean cell. Nighttime light was quantified using the global annual simulated VIIRS Nighttime Light dataset (SVNL) (42), which provides a continuous, sensor-harmonized record from 1992 to 2023 at 500-m resolution. The SVNL series is produced by cross-calibrating DMSP-OLS and VIIRS data with a U-Net framework to yield annually consistent VIIRS-like radiance values. For consistency with other environmental datasets, we used SVNL data only for the same 1993–2020 period. For each site and year, we extracted the mean annual radiance within a 5-km radius buffer using QGIS v3.36 and used these values in downstream analyses.

From the monthly layers we derived, for each population and year, the following summaries: mean 2-m air temperature and its annual range, mean sea-surface temperature, vector-averaged 10-m wind speed (annual mean and SD), annual precipitation total (converting ERA5-Land hourly precipitation rates to mm per month and summing over months) and its seasonality (coefficient of variation, %), mean sea salinity and its annual range, and a vegetation index defined as high minus low fractional cover. Nighttime light values were log-transformed (log1p) prior to analysis. Data handling was performed with *terra* and *sf* packages in R v4.4.2.

To identify environmental predictors associated with species occurrence, we fitted a Bayesian logistic regression with species (*L. furcata* vs. *L. laticarpa*) as the binary response (*L. laticarpa* coded as 1), z-scored environmental variables (listed above) as fixed effects, and random effects for sampling site and year. The model included both random intercepts and a random slope for nighttime light intensity by site to account for spatially varying sensitivity to illumination. Models were implemented in brms using a horseshoe prior on fixed effects (to promote sparsity) and weakly informative Student-*t* and exponential priors on the intercept and random-effect standard deviations, respectively. Four independent Markov chains were run for 4,000 iterations each with stringent sampler controls (high adapt_delta and increased tree depth) to ensure convergence. Posterior means and 95% credible intervals were summarized for each predictor, and the directionality of effects (probability of being above or below zero) was quantified. Predictors were ranked by the absolute posterior mean, and posterior densities as well as violin and boxplot summaries of raw predictor values by species were visualized (Fig. S3).

### Rearing conditions

Based on the sequencing results, we selected three populations for each of *L. furcata* and *L. laticarpa* near the boundary of their distributions (toy, ryu, and tsh for *L. furcata*; onu, fut, and sod for *L. laticarpa*; highlighted with black-rimmed circles in Fig. 1a). These populations were chosen for subsequent experiments aimed at examining traits associated with ecological divergence, while avoiding potential confounding effects such as genetic introgression from non-native *Ligia* species that occurs in heavily urbanized port areas near Tokyo as shown in Fig. 1a. From each population, we collected approximately 40–60 juvenile (1-cm-long) *Ligia* individuals in July 2024 for rearing and behavioral analyses.

Juvenile *Ligia* collected from each site were randomly assigned to one of two nighttime light treatments for rearing: one with ALAN and one without. The air temperature was maintained at 23°C under a 12-hour light/12-hour dark cycle for both treatments, with lights turned on at 06:00 and off at 18:00. During the dark period, light intensity was approximately 10 lx in the ALAN treatment and 0 lx in the control (43). Daytime light intensity was kept at around 2500 lx, corresponding to an overcast day in the wild (5).

Groups of *Ligia* individuals were reared in polypropylene containers measuring 9.2 cm in width, 27.8 cm in length and 18 cm in height and were fed weekly with Hikari large turtle pellets (Kyorin Co. Ltd., Japan). A 2-mm mesh made of the same polypropylene material was placed at the bottom of each container. In total, 12 rearing containers were prepared, corresponding to 6 populations × 2 rearing conditions (with or without ALAN), and these were distributed evenly across four plastic storage boxes. Three containers were inclined and placed on a PVC pipe frame within each storage box (44 cm wide × 74 cm deep × 35 cm high). The boxes were filled with seawater (salinity ca. 1.023 g/cm^3^) to a depth of approximately 15 cm and equipped with a filtration system for water circulation. A schematic illustration of the rearing setup is provided in Fig. S4. This configuration created both aquatic and terrestrial zones within the tilted containers, enabling suitable conditions for *Ligia*. To eliminate potential positional effects within or between storage boxes, the rearing containers were randomly rearranged every 3–7 days during survival monitoring (see below).

### Evaluation of survival and growth

To evaluate survival under different light conditions, the number of live individuals in each container was recorded approximately every 3 to 7 days over a period of five months. Dead individuals were removed promptly upon detection. Survival curves were generated for each group and analyzed using a Cox proportional hazards regression model to test for the effects of species and rearing condition. Body length was measured at the start of rearing and after one month as an indicator of growth, and again at five months during the activity rhythm experiments. To assess body size, individuals were photographed from above alongside a ruler, either collectively in a tray (for measurements at the start and after one month of rearing) or individually in tubes (at five months, during the activity rhythm experiments).

Body length was defined as the distance from the anterior margin of the head to the posterior end of the abdomen and was measured using ImageJ v2.14.0. Measurements were calibrated using the scale provided by the ruler in each image.

### Evaluation of activity rhythms

At five months after the start of rearing, in December 2024, we investigated the effects of chronic exposure to ALAN on the activity rhythms of *Ligia*. Daily activity was monitored under the 12L:12D cycle using two Locomotor Activity Monitors (LAM25; Trikinetics Inc., Waltham, MA, USA), each capable of simultaneously measuring 32 individuals, allowing for the activity of 64 individuals to be recorded per experimental day. Two to fifteen individuals per group were used for activity measurements, depending on survival outcomes in each population and treatment condition. Each *Ligia* was individually placed into a transparent tube (25 mm in diameter and 100 mm in length), sealed at both ends with cotton soaked in seawater to ensure proper respiration. To eliminate potential positional effects within the activity monitor, all individuals were randomly assigned to tube positions, regardless of species, population, or rearing condition. Locomotor activity was recorded via infrared sensors within the LAM, and the accumulated number of times each individual crossed the beam was automatically counted every 15 seconds by DAMSystem3 Software (Trikinetics Inc.). Each trial involved continuous monitoring over a single day—from early afternoon until the following evening. The experiment consisted of four sessions: sessions 1 and 4 were conducted with ALAN during the night, while sessions 2 and 3 were conducted in complete darkness. Each individual was tested twice—either in sessions 1 and 3 or in sessions 2 and 4—under both ALAN and control conditions, with a three-day interval between the two trials. Trial order was randomized across subgroups to control for order effects.

One month later, in January 2025, to assess the endogenous rhythms of *Ligia*, we conducted an additional experiment in which individuals were placed in complete darkness for four consecutive days. In this experiment, each *Ligia* was housed in a transparent acrylic case (56 mm in width, 80 mm in length and 50 mm in height) with a transparent bottom and opaque white side walls. The white walls were used to minimize visual stimuli from the external environment, including the movement of other individuals. A 3-mm thick vinyl perforated board was placed on the bottom to facilitate movement, and an appropriate volume of seawater was added to ensure proper respiration. A total of 15 cases were recorded simultaneously per session, and the experiment was repeated eight times to obtain sufficient data across individuals. One trial (28 January 2025) was excluded due to lid misplacement that caused excessive mortality. To eliminate positional effects, the spatial arrangement of individuals within each session was randomized. An infrared light board was installed beneath the case, and an infrared camera (DMK33UX290, The Imaging Source, Germany) positioned above captured images at one-minute intervals over the four-day period.

Individual movements were quantified from the time-lapse images using a custom Python script. Circadian rhythms were analyzed based on activity time series using the ac_periodogram() function implemented in the *zeitgebr* package in R.

### Statistical analysis

All statistical analyses were conducted using R v.4.4.2. Wilcoxon’s rank sum test was used to estimate differences in environmental variables and mean activity between species. The Cox proportional hazards model was used to evaluate the effects of rearing conditions and species on survival. Wilcoxon’s signed rank test was used to compare mean activity levels between night and light-on periods for each rearing and experimental condition. Linear mixed models (LMMs) were used to assess the effects of ALAN treatment and species on body length, mean daily activity, and endogenous circadian rhythms. Population, LAM ID, tube or case position, experimental date, and trial order (where applicable) were included as random effects. The significance of fixed effects in the LMMs and the Cox model was tested using Type II Wald *χ*^2^ test.

## Acknowledgement

We are grateful to Dr. Yuma Takahashi for valuable comments on the behavioral assays and on the manuscript draft. We also thank Dr. Mitsuhiko P. Sato for helpful advice on the genomic analyses.

## Fundings

This work was supported by the Japan Society for the Promotion of Science (Grants-in-Aid for Scientific Research 25K18517) and Obayashi Foundation.

## Competing interests

The author declares no competing interests.

## Data and code availability

Data and codes to reproduce the findings described in the present study will be available upon publication.

## Notes

### Competing Interest Statement

The authors have declared no competing interest.

