## Supplementary Information for "Night light of a metropolitan coastline aligns with habitat segregation and divergent plasticity among closely related coastal isopods"

This file includes:

- **Tables S1–5**
- **Figure S1–6**

**Table S1. Information of sequenced samples.**

| Sample ID | Sampling site | Longitude (°N) | Latitude (°E) | Inferred species | Genbank ID (16S rDNA) | Genbank ID (18S rDNA) | SRA ID (MIG-seq) |
|---|---|---|---|---|---|---|---|
| ito1 | Tateyama city, Chiba | 34.9572 | 139.7687 | *L. furcata* | Pending | Pending | SRR35820454 |
| ito2 | | | | *L. furcata* | Pending | Pending | SRR35820453 |
| ito3 | | | | *L. furcata* | – | – | SRR35820330 |
| ito4 | | | | *L. furcata* | – | – | SRR35820415 |
| ito5 | | | | *L. furcata* | – | – | SRR35820404 |
| ito6 | | | | *L. furcata* | – | – | SRR35820393 |
| ito7 | | | | *L. furcata* | – | – | SRR35820318 |
| ito8 | | | | *L. furcata* | – | – | SRR35820307 |
| haz1 | Tateyama city, Chiba | 34.9749 | 139.7824 | *L. furcata* | Pending | Pending | – |
| haz2 | | | | *L. furcata* | Pending | Pending | – |
| oki1 | Tateyama city, Chiba | 34.9911 | 139.8370 | *L. furcata* | Pending | Pending | SRR35820296 |
| oki2 | | | | *L. furcata* | Pending | Pending | SRR35820285 |
| oki3 | | | | *L. furcata* | – | – | SRR35820452 |
| oki4 | | | | *L. furcata* | – | – | SRR35820441 |
| oki5 | | | | *L. furcata* | – | – | SRR35820430 |
| oki6 | | | | *L. furcata* | – | – | SRR35820419 |
| oki7 | | | | *L. furcata* | – | – | SRR35820376 |
| oki8 | | | | *L. furcata* | – | – | SRR35820365 |
| toy1 | Minamiboso city, Chiba | 35.0521 | 139.8312 | *L. furcata* | Pending | Pending | SRR35820354 |
| toy2 | | | | *L. furcata* | Pending | Pending | SRR35820343 |
| toy3 | | | | *L. furcata* | – | – | SRR35820332 |
| toy4 | | | | *L. furcata* | – | – | SRR35820331 |
| toy5 | | | | *L. furcata* | – | – | SRR35820329 |
| toy6 | | | | *L. furcata* | – | – | SRR35820328 |
| toy7 | | | | *L. furcata* | – | – | SRR35820327 |

| | | | | | | | |
|---|---|---|---|---|---|---|---|
| toy8 | | | | *L. furcata* | – | – | SRR35820326 |
| iwa1 | Minamiboso city, Chiba | 35.0871 | 139.8428 | *L. furcata* | Pending | Pending | – |
| iwa2 | | | | *L. furcata* | Pending | Pending | – |
| ryu1 | Kyonan town, Chiba | 35.1185 | 139.8282 | *L. furcata* | Pending | Pending | SRR35820325 |
| ryu2 | | | | *L. furcata* | Pending | Pending | SRR35820324 |
| ryu3 | | | | *L. furcata* | – | – | SRR35820323 |
| ryu4 | | | | *L. furcata* | – | – | SRR35820418 |
| ryu5 | | | | *L. furcata* | – | – | SRR35820417 |
| ryu6 | | | | *L. furcata* | – | – | SRR35820416 |
| ryu7 | | | | *L. furcata* | – | – | SRR35820414 |
| ryu8 | | | | *L. furcata* | – | – | SRR35820413 |
| mot1 | Kyonan town, Chiba | 35.1451 | 139.8305 | *L. furcata* | Pending | Pending | – |
| mot2 | | | | *L. furcata* | Pending | Pending | – |
| tsh1 | Futtsu city, Chiba | 35.2050 | 139.8358 | *L. furcata* | Pending | Pending | SRR35820412 |
| tsh2 | | | | *L. furcata* | Pending | Pending | SRR35820411 |
| tsh3 | | | | *L. furcata* | – | – | SRR35820410 |
| tsh4 | | | | *L. furcata* | – | – | SRR35820409 |
| tsh5 | | | | *L. furcata* | – | – | SRR35820408 |
| tsh6 | | | | *L. furcata* | – | – | SRR35820407 |
| tsh7 | | | | *L. furcata* | – | – | SRR35820406 |
| tsh8 | | | | *L. furcata* | – | – | SRR35820405 |
| onu1 | Futtsu city, Chiba | 35.2784 | 139.8476 | *L. laticarpa* | Pending | Pending | SRR35820403 |
| onu2 | | | | *L. laticarpa* | Pending | Pending | SRR35820402 |
| onu3 | | | | *L. laticarpa* | – | – | SRR35820401 |
| onu4 | | | | *L. laticarpa* | – | – | SRR35820400 |
| onu5 | | | | *L. laticarpa* | – | – | SRR35820399 |
| onu6 | | | | *L. laticarpa* | – | – | SRR35820398 |
| onu7 | | | | *L. laticarpa* | – | – | SRR35820397 |
| onu8 | | | | *L. laticarpa* | – | – | SRR35820396 |

| | | | | | | | |
|---|---|---|---|---|---|---|---|
| fut1 | Futtsu city, Chiba | 35.3272 | 139.8218 | *L. laticarpa* | Pending | Pending | SRR35820395 |
| fut2 | | | | *L. laticarpa* | Pending | Pending | SRR35820394 |
| fut3 | | | | *L. laticarpa* | – | – | SRR35820392 |
| fut4 | | | | *L. laticarpa* | – | – | SRR35820391 |
| fut5 | | | | *L. laticarpa* | – | – | SRR35820390 |
| fut6 | | | | *L. laticarpa* | – | – | SRR35820389 |
| fut7 | | | | *L. laticarpa* | – | – | SRR35820388 |
| fut8 | | | | *L. laticarpa* | – | – | SRR35820387 |
| tor1 | Kisarazu city, Chiba | 35.3826 | 139.9128 | *L. laticarpa* | Pending | Pending | SRR35820322 |
| tor2 | | | | *L. laticarpa* | Pending | Pending | SRR35820321 |
| tor3 | | | | *L. laticarpa* | – | – | SRR35820320 |
| tor4 | | | | *L. laticarpa* | – | – | SRR35820319 |
| tor5 | | | | *L. laticarpa* | – | – | SRR35820317 |
| tor6 | | | | *L. laticarpa* | – | – | SRR35820316 |
| tor7 | | | | *L. laticarpa* | – | – | SRR35820315 |
| tor8 | | | | *L. laticarpa* | – | – | SRR35820314 |
| sod1 | Sodegaura city, Chiba | 35.4537 | 139.9507 | *L. laticarpa* | Pending | Pending | SRR35820313 |
| sod2 | | | | *L. laticarpa* | Pending | Pending | SRR35820312 |
| sod3 | | | | *L. laticarpa* | – | – | SRR35820311 |
| sod4 | | | | *L. laticarpa* | – | – | SRR35820310 |
| sod5 | | | | *L. laticarpa* | – | – | SRR35820309 |
| sod6 | | | | *L. laticarpa* | – | – | SRR35820308 |
| sod7 | | | | *L. laticarpa* | – | – | SRR35820306 |
| sod8 | | | | *L. laticarpa* | – | – | SRR35820305 |
| ori1 | Ichihara city, Chiba | 35.5427 | 140.0604 | *L. laticarpa* | Pending | Pending | SRR35820304 |
| ori2 | | | | *L. laticarpa* | Pending | Pending | SRR35820303 |
| ori3 | | | | *L. laticarpa* | – | – | SRR35820302 |
| ori4 | | | | *L. laticarpa* | – | – | SRR35820301 |
| ori5 | | | | *L. laticarpa* | – | – | SRR35820300 |

| | | | | | | | |
|---|---|---|---|---|---|---|---|
| ori6 | | | | *L. laticarpa* | – | – | SRR35820299 |
| ori7 | | | | *L. laticarpa* | – | – | SRR35820298 |
| ori8 | | | | *L. laticarpa* | – | – | SRR35820297 |
| ina1 | Chiba city, Chiba | 35.6122 | 140.0678 | *L. laticarpa* | Pending | Pending | SRR35820295 |
| ina2 | | | | *L. laticarpa* | Pending | Pending | SRR35820294 |
| ina3 | | | | *L. laticarpa* | – | – | SRR35820293 |
| ina4 | | | | *L. laticarpa* | – | – | SRR35820292 |
| ina5 | | | | *L. laticarpa* | – | – | SRR35820291 |
| ina6 | | | | *L. laticarpa* | – | – | SRR35820290 |
| ina7 | | | | *L. laticarpa* | – | – | SRR35820289 |
| ina8 | | | | *L. laticarpa* | – | – | SRR35820288 |
| san1 | Funabashi city, Chiba | 35.6704 | 139.9713 | *L. cinerascens* | Pending | Pending | SRR35820287 |
| san2 | | | | *L. laticarpa* | Pending | Pending | SRR35820286 |
| san3 | | | | *L. cinerascens* | – | – | SRR35820284 |
| san4 | | | | *L. laticarpa* | – | – | SRR35820283 |
| san5 | | | | *L. laticarpa* | – | – | SRR35820282 |
| san6 | | | | *L. laticarpa* | – | – | SRR35820281 |
| san7 | | | | *L. laticarpa* | – | – | SRR35820280 |
| san8 | | | | *L. laticarpa* | – | – | SRR35820279 |
| nag1 | Edogawa ward, Tokyo | 35.6474 | 139.8832 | *L. laticarpa* | Pending | Pending | SRR35820278 |
| nag2 | | | | *L. laticarpa* | Pending | Pending | SRR35820277 |
| nag3 | | | | *L. laticarpa* | – | – | SRR35820276 |
| nag4 | | | | *L. laticarpa* | – | – | SRR35820275 |
| nag5 | | | | *L. laticarpa* | – | – | SRR35820451 |
| nag6 | | | | *L. laticarpa* | – | – | SRR35820450 |
| nag7 | | | | *L. laticarpa* | – | – | SRR35820449 |
| nag8 | | | | *L. laticarpa* | – | – | SRR35820448 |
| tsb1 | Ohta ward, Tokyo | 35.5804 | 139.7870 | *L. laticarpa* | Pending | Pending | SRR35820447 |
| tsb2 | | | | *L. laticarpa* | Pending | Pending | SRR35820446 |

| | | | | | | | |
|---|---|---|---|---|---|---|---|
| tsb3 | | | | *L. laticarpa* | – | – | SRR35820445 |
| tsb4 | | | | *L. laticarpa* | – | – | SRR35820444 |
| tsb5 | | | | *L. laticarpa* | – | – | SRR35820443 |
| tsb6 | | | | *L. laticarpa* | – | – | SRR35820442 |
| tsb7 | | | | *L. laticarpa* | – | – | SRR35820440 |
| tsb8 | | | | *L. laticarpa* | – | – | SRR35820439 |
| tok1 | Ohta ward, Tokyo | 35.5772 | 139.7568 | *L. cinerascens* | Pending | Pending | SRR35820345 |
| tok2 | | | | *L. cinerascens* | Pending | Pending | SRR35820344 |
| tok3 | | | | *L. cinerascens* | – | – | SRR35820342 |
| tok4 | | | | *L. cinerascens* | – | – | SRR35820341 |
| tok5 | | | | *L. cinerascens* | – | – | SRR35820340 |
| tok6 | | | | *L. cinerascens* | – | – | SRR35820339 |
| tok7 | | | | *L. cinerascens* | – | – | SRR35820338 |
| tok8 | | | | *L. cinerascens* | – | – | SRR35820337 |
| kaw1 | Kawasaki city, Kanagawa | 35.5051 | 139.7756 | *L. laticarpa* | Pending | Pending | SRR35820438 |
| kaw2 | | | | *L. laticarpa* | Pending | Pending | SRR35820437 |
| kaw3 | | | | *L. cinerascens* | – | – | SRR35820436 |
| kaw4 | | | | *L. laticarpa* | – | – | SRR35820435 |
| kaw5 | | | | *L. cinerascens* | – | – | SRR35820434 |
| kaw6 | | | | *L. cinerascens* | – | – | SRR35820433 |
| kaw7 | | | | *L. laticarpa* | – | – | SRR35820432 |
| kaw8 | | | | *L. laticarpa* | – | – | SRR35820431 |
| dai1 | Yokohama city, Kanagawa | 35.4510 | 139.6971 | *L. laticarpa* | Pending | Pending | SRR35820429 |
| dai2 | | | | *L. laticarpa* | Pending | Pending | SRR35820428 |
| dai3 | | | | *L. laticarpa* | – | – | SRR35820427 |
| dai4 | | | | *L. laticarpa* | – | – | SRR35820426 |
| dai5 | | | | *L. laticarpa* | – | – | SRR35820425 |
| dai6 | | | | *L. laticarpa* | – | – | SRR35820424 |
| dai7 | | | | *L. laticarpa* | – | – | SRR35820423 |

| | | | | | | | |
|---|---|---|---|---|---|---|---|
| dai8 | | | | *L. laticarpa* | – | – | SRR35820422 |
| hak1 | Yokohama city, Kanagawa | 35.3404 | 139.6398 | *L. laticarpa* | Pending | Pending | SRR35820421 |
| hak2 | | | | *L. laticarpa* | Pending | Pending | SRR35820420 |
| hak3 | | | | *L. laticarpa* | – | – | SRR35820386 |
| hak4 | | | | *L. laticarpa* | – | – | SRR35820385 |
| hak5 | | | | *L. laticarpa* | – | – | SRR35820384 |
| hak6 | | | | *L. laticarpa* | – | – | SRR35820383 |
| hak7 | | | | *L. laticarpa* | – | – | SRR35820382 |
| hak8 | | | | *L. laticarpa* | – | – | SRR35820381 |
| yok1 | Yokosuka city, Kanagawa | 35.2769 | 139.6837 | *L. laticarpa* | Pending | Pending | SRR35820380 |
| yok2 | | | | *L. laticarpa* | Pending | Pending | SRR35820379 |
| yok3 | | | | *L. laticarpa* | – | – | SRR35820378 |
| yok4 | | | | *L. laticarpa* | – | – | SRR35820377 |
| yok5 | | | | *L. laticarpa* | – | – | SRR35820375 |
| yok6 | | | | *L. laticarpa* | – | – | SRR35820374 |
| yok7 | | | | *L. laticarpa* | – | – | SRR35820373 |
| yok8 | | | | *L. laticarpa* | – | – | SRR35820372 |
| tom1 | Yokosuka city, Kanagawa | 35.2331 | 139.7291 | *L. laticarpa* | Pending | Pending | SRR35820371 |
| tom2 | | | | *L. laticarpa* | Pending | Pending | SRR35820370 |
| tom3 | | | | *L. laticarpa* | – | – | SRR35820369 |
| tom4 | | | | *L. laticarpa* | – | – | SRR35820368 |
| tom5 | | | | *L. laticarpa* | – | – | SRR35820367 |
| tom6 | | | | *L. laticarpa* | – | – | SRR35820366 |
| tom7 | | | | *L. laticarpa* | – | – | SRR35820364 |
| tom8 | | | | *L. laticarpa* | – | – | SRR35820363 |
| miu1 | Yokosuka city, Kanagawa | 35.1965 | 139.6684 | *L. furcata* | Pending | Pending | SRR35820362 |
| miu2 | | | | *L. furcata* | Pending | Pending | SRR35820361 |
| miu3 | | | | *L. furcata* | – | – | SRR35820360 |
| miu4 | | | | *L. furcata* | – | – | SRR35820359 |

| | | | | | | | |
|---|---|---|---|---|---|---|---|
| miu5 | | | | *L. furcata* | – | – | SRR35820358 |
| miu6 | | | | *L. furcata* | – | – | SRR35820357 |
| miu7 | | | | *L. furcata* | – | – | SRR35820356 |
| miu8 | | | | *L. furcata* | – | – | SRR35820355 |
| mag1 | Miura city, Kanagawa | 35.1453 | 139.6770 | *L. furcata* | Pending | Pending | – |
| mag2 | | | | *L. furcata* | Pending | Pending | – |
| mis1 | Miura city, Kanagawa | 35.1368 | 139.6118 | *L. furcata* | Pending | Pending | SRR35820354 |
| mis2 | | | | *L. furcata* | Pending | Pending | SRR35820353 |
| mis3 | | | | *L. furcata* | – | – | SRR35820352 |
| mis4 | | | | *L. furcata* | – | – | SRR35820351 |
| mis5 | | | | *L. furcata* | – | – | SRR35820350 |
| mis6 | | | | *L. furcata* | – | – | SRR35820349 |
| mis7 | | | | *L. furcata* | – | – | SRR35820348 |
| mis8 | | | | *L. furcata* | – | – | SRR35820347 |
| kat1 | Kushiro city, Hokkaido | 42.9454 | 144.4463 | *L. cinerascens* | Pending | Pending | SRR35820336 |
| kat2 | | | | *L. cinerascens* | Pending | Pending | SRR35820335 |
| kat3 | | | | *L. cinerascens* | – | – | SRR35820334 |
| kat4 | | | | *L. cinerascens* | – | – | SRR35820333 |

**Table S2. Posterior effects of environmental variables distinguishing *L. furcata* and *L. laticarpa*.** Response coded as *L. laticarpa* = 1 and *L. furcata* = 0; predictors are z-scored; site and year included as a random intercept. Reported are posterior mean, 95% credible interval (CI), probability of direction ($p_d$). Positive coefficients indicate higher probability of *L. laticarpa* occurrence.

| Item | Posterior mean | 95% CI upper | 95% CI lower | $p_d$ |
|---|---:|---:|---:|---:|
| Salinity (mean) | –7.009 | –16.158 | –0.252 | 0.983 |
| Nighttime light | 4.150 | –0.034 | 9.738 | 0.966 |
| Vegetation index | –4.121 | –12.120 | 0.158 | 0.942 |
| Sea surface temperature (mean) | –1.655 | –6.813 | 0.978 | 0.789 |
| Salinity (range) | 1.568 | –0.964 | 6.485 | 0.782 |
| Land temperature (mean) | 1.400 | –0.634 | 5.511 | 0.805 |
| Annual precipitation (CV) | 0.947 | –0.368 | 3.496 | 0.843 |
| Wind speed (mean) | –0.583 | –3.375 | 1.252 | 0.694 |
| Wind speed (SD) | –0.581 | –2.994 | 1.050 | 0.720 |
| Land temperature (range) | 0.243 | –1.982 | 2.906 | 0.587 |
| Annual precipitation | –0.012 | –1.741 | 1.875 | 0.511 |

**Table S3. Effect of species, rearing condition, and days since the start of experiment on body length of *Ligia*, estimated by LMM.** Population was treated as a random effect.

| Item | $\chi^2$ | df | $P$ |
|---|---|---|---|
| Species | 0.778 | 1 | 0.378 |
| Rearing condition | 1.622 | 1 | 0.203 |
| Rearing duration | 1797.744 | 2 | $< 2.0\times10^{-16}$ |
| Species × Rearing condtion | 5.178 | 1 | $2.288\times10^{-2}$ |
| Species × Rearing duration | 34.135 | 2 | $5.141\times10^{-9}$ |
| Rearing condition × Rearing duration | 7.904 | 2 | $4.932\times10^{-3}$ |
| Species × Rearing condition × Rearing duration | 0.404 | 2 | 0.525 |

**Table S4. Effect of species, rearing condition, experimental condition, and body length on mean total activity in a day, estimated by LMM.** Population, LAM ID, tube position, experimental date and experimental order were treated as a random effect.

| Item | χ2 | df | *P* |
|---|---|---|---|
| Species | 4.497 | 1 | 0.034 |
| Rearing condition | 4.520 | 1 | 0.034 |
| Experimental condition | 0.996 | 1 | 0.318 |
| Body length | 46.031 | 1 | $1.161 \times 10^{-11}$ |
| Species × Rearing condition | 7.232 | 1 | 0.007 |
| Species × Experimental condition | 0.000 | 1 | 0.998 |
| Species × Body length | 0.159 | 1 | 0.690 |
| Rearing condtion × Experimental condition | 1.942 | 1 | 0.163 |
| Rearing condtion × Body length | 1.407 | 1 | 0.236 |
| Experimental condtion × Body length | 0.140 | 1 | 0.709 |
| Species × Rearing condition × Experimental condition | 1.179 | 1 | 0.278 |
| Species × Rearing condition × Body length | 5.103 | 1 | 0.024 |
| Species × Experimental condition × Body length | 0.068 | 1 | 0.795 |
| Rearing condition × Experimental condition × Body length | 4.557 | 1 | 0.033 |
| Species × Rearing condition × Experimental condition × Body length | 0.197 | 1 | 0.658 |

**Table S5. Effect of species and rearing condition on endogenous circadian rhythms, estimated by LMM.** Population, case position and experimental date were treated as a random effect.

| Item | $\chi^2$ | df | *P* |
|---|---|---|---|
| Species | 0.461 | 1 | 0.497 |
| Rearing condition | 0.009 | 1 | 0.926 |
| Species × Rearing condtion | 4.241 | 1 | 0.039 |

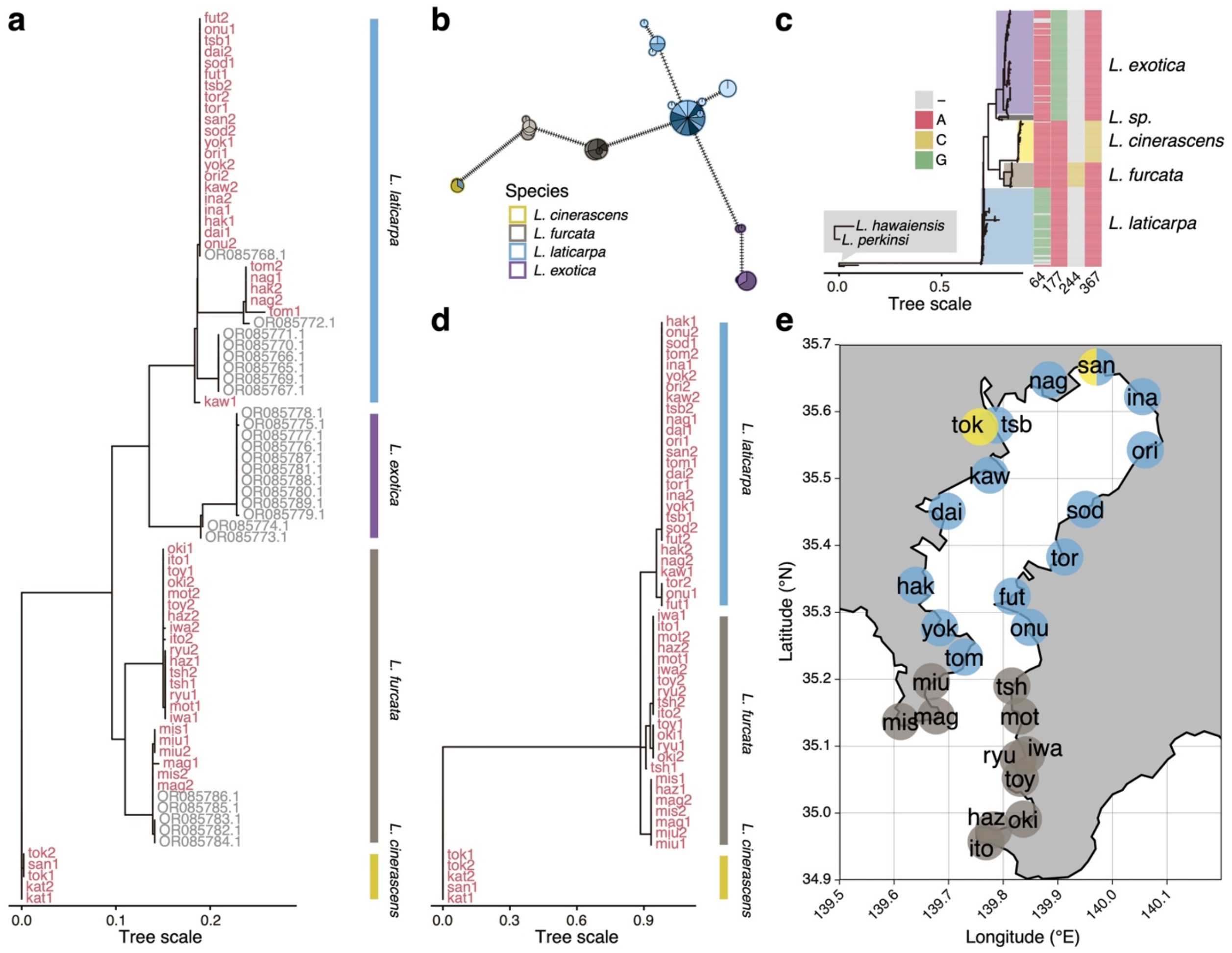


**Figure S1. Phylogenetic and geographic differentiation of *Ligia* species along the Tokyo Bay coastline.**

**(a)** Maximum-likelihood tree based on mitochondrial 16S rRNA gene sequences showing major clades corresponding to *L. laticarpa*, *L. exotica*, *L. furcata*, and *L. cinerascens*. Newly sequenced samples from this study are shown in red, with reference sequences (1) indicated in gray. **(b)** Median-joining haplotype network of the 16S sequences illustrating relationships among haplotypes and species clusters. The inner color of each pie indicates the sampling site, while the outline color represents the inferred species. **(c)** Full alignment of 16S sequences with 229 previously sequenced *Ligia* samples (1–7) revealed diagnostic nucleotide substitutions distinguishing the four species, with positions 64, 177, 244, and 367 highlighted (see Supplementary Data 1 for the full sequence alignment). A yet undescribed *Ligia* species closely related to *L. exotica* (*L. sp.* from Okinawa; 1) was provisionally treated as *L. exotica* for consistency. **(d)** Maximum-likelihood tree based on nuclear 18S rRNA gene sequences, confirming species-level groupings consistent with the mitochondrial tree. **(e)** Geographic distribution of *Ligia* populations analyzed around Tokyo Bay. Sampling sites are colored by species inferred by the target sequencing: *L. laticarpa* (blue), *L. furcata* (brown), and *L. cinerascens* (yellow). Population codes correspond to those in Table S1.

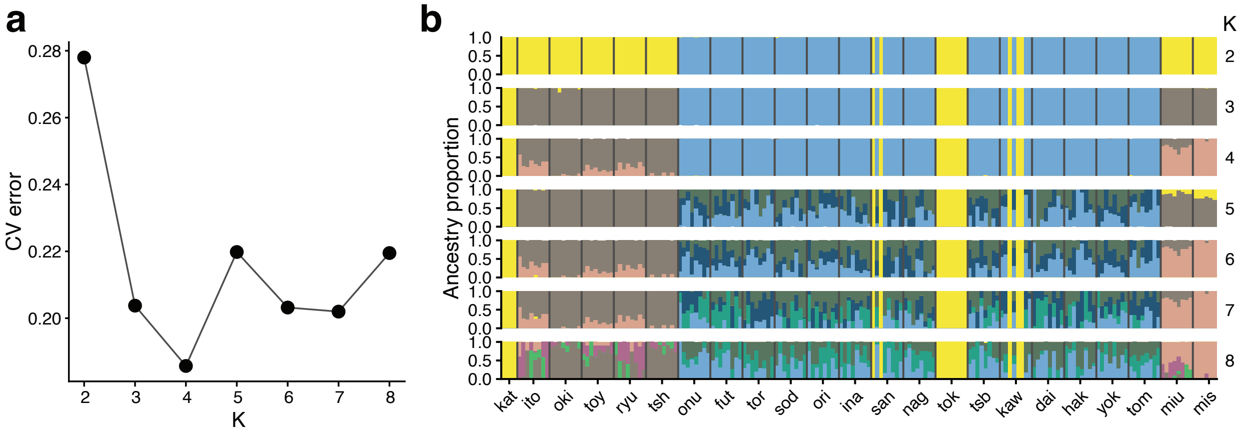


**Figure S2. Population structure inferred with ADMIXTURE.**

**(a)** Cross-validation (CV) error across $K = 2–8$. The x-axis shows K and the y-axis the CV error reported by ADMIXTURE; the minimum occurs at $K = 4$. **(b)** Individual ancestry proportions (Q-values) for $K = 2–8$ shown as stacked bars. Vertical gray lines indicate population boundaries; bottom labels are population codes. Colors denote genetic clusters.

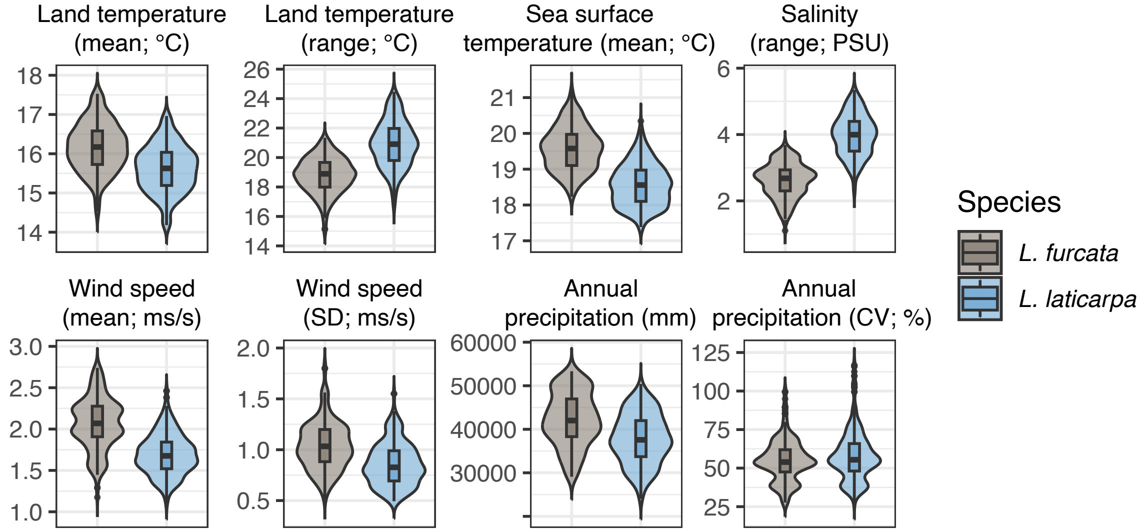


**Figure S3. Environmental differences between habitats of *L. furcata* and *L. laticarpa*.** Violin plots and boxplots showing species-wise comparisons of environmental variables across sampling sites.

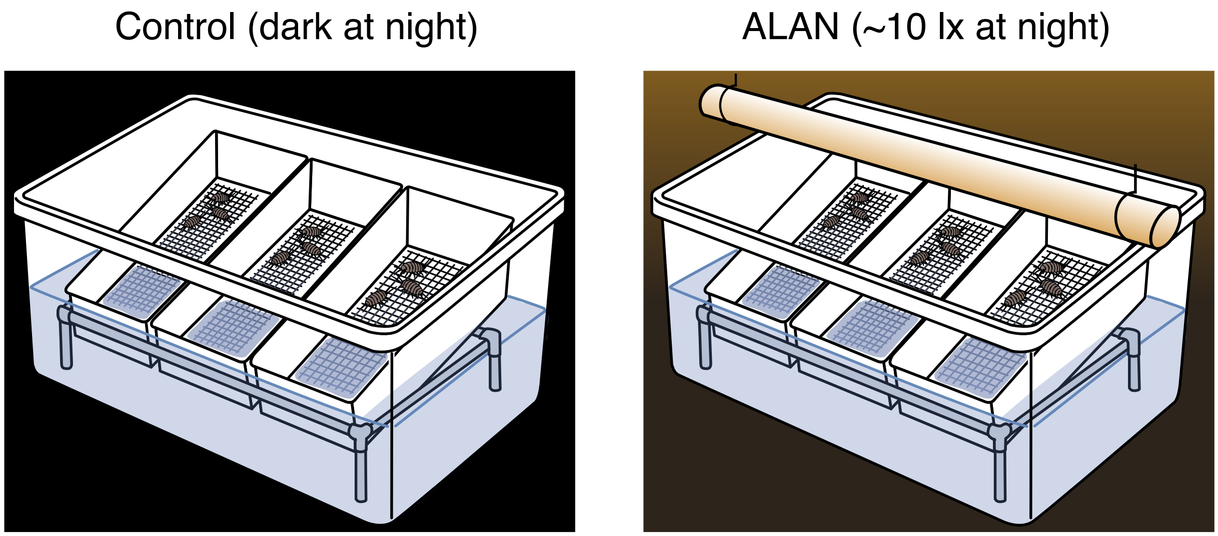


**Figure S4. Experimental setup for control and ALAN treatments.**

Juvenile *Ligia* were reared at 23°C under a 12-hour light/12-hour dark cycle. During the dark phase, individuals were either kept in complete darkness (control; left panel) or exposed to dim artificial light at ~10 lx (ALAN; right panel). Each rearing unit consisted of three tilted polypropylene containers placed on a PVC pipe frame within a plastic storage box filled with, providing both aquatic and terrestrial environments. A total of 12 rearing containers (6 populations × 2 lighting conditions) were evenly distributed across four storage boxes. Containers were randomly repositioned every 3–7 days to minimize potential positional effects.

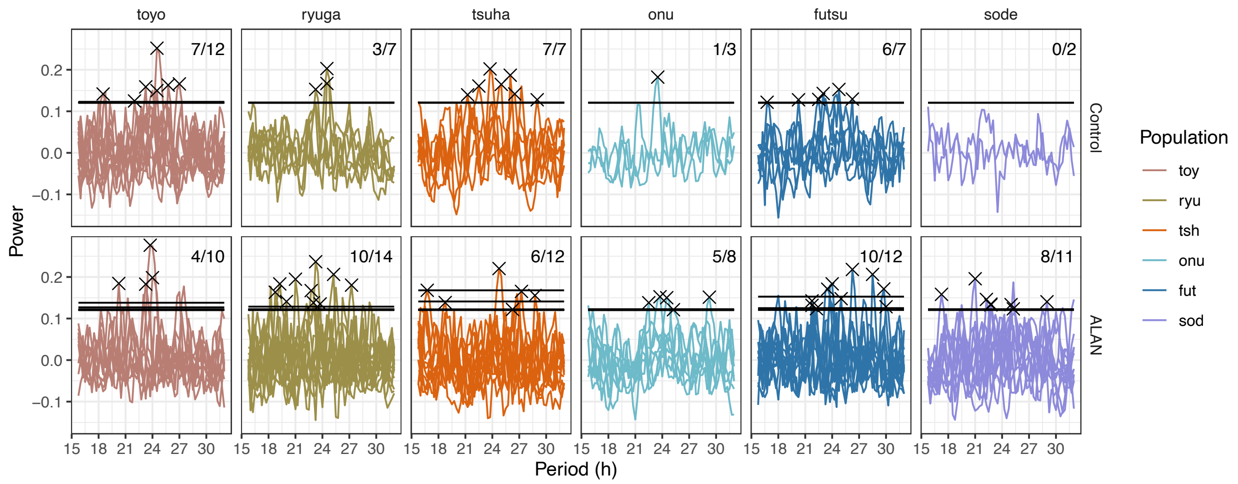


**Figure S5. Periodogram analysis of circadian rhythmicity under constant darkness.** Autocorrelation-based periodograms for each population under control and ALAN rearing conditions. Black '×' marks indicate significant peaks. Horizontal black lines denote the significance thresholds. Differences in rhythm strength and period distributions suggest species- and condition-dependent effects of ALAN exposure on endogenous activity rhythms. The upper-right corner of each panel shows the number of individuals exhibiting significant rhythmic peaks out of the total number analysed for that condition.

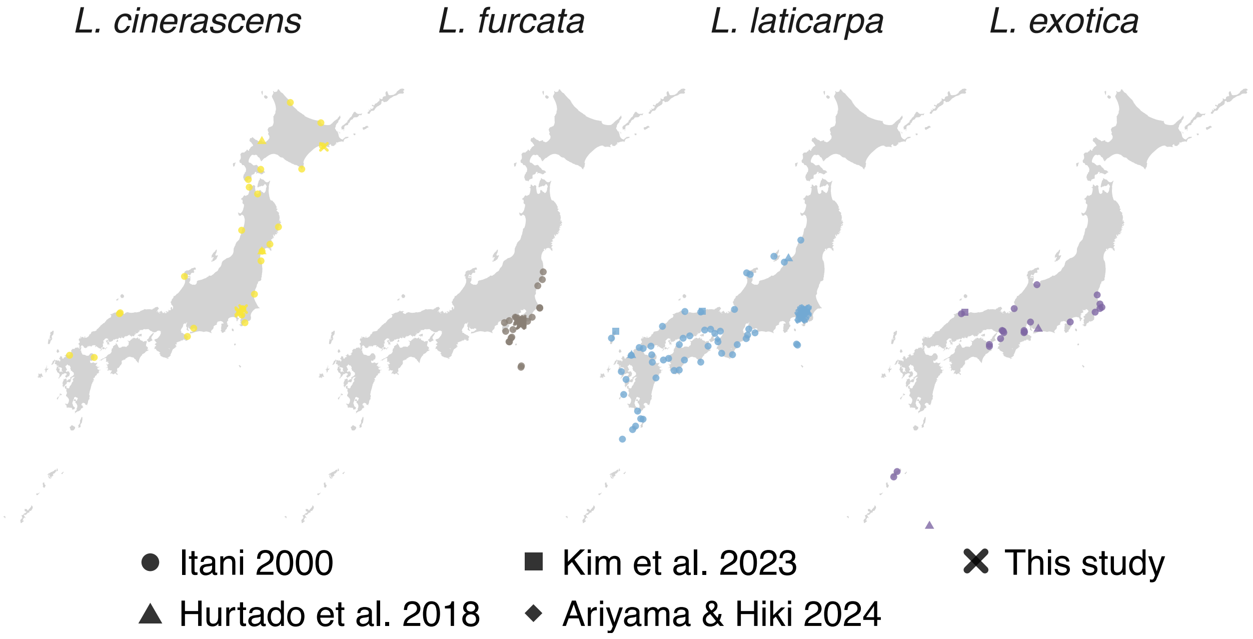


**Figure S6. Nationwide distribution of coastal *Ligia* species in Japan based on prior molecular studies and the present work.**

Geographic ranges of four *Ligia* species (*L. cinerascens*, *L. furcata*, *L. laticarpa*, and *L. exotica*) compiled from published molecular studies (1, 5, 6, 8) and from this study. Colored points mark occurrence localities assigned to species by mitochondrial clades (COI or 16S): yellow = *L. cinerascens*, brown = *L. furcata*, blue = *L. laticarpa*, purple = *L. exotica*. Point shape denotes the source study.
